# DNA binding activity of CAMTA3 is essential for its function: Identification of critical amino acids for its transcriptional activity

**DOI:** 10.1101/2023.06.22.546133

**Authors:** Kasavajhala V.S.K. Prasad, Amira Abdel-Hameed, Qiyan Jiang, Anireddy S.N. Reddy

## Abstract

Calmodulin-binding transcription activators (CAMTAs), a small family of highly conserved transcription factors, function in calcium-mediated signaling pathways. Of the six CAMTAs in Arabidopsis, CAMTA3 (also referred to as SR1) regulates diverse biotic and abiotic stress responses. A recent study has shown that CAMTA3 is a guardee of NLR ((nucleotide-binding, leucine-rich repeat domain) immune receptors in modulating plant immunity, thereby suggesting that CAMTA3 transcriptional activity is not necessary for its function. Here, we show that the DNA-binding activity of CAMTA3 is essential for its role in mediating plant immune responses. Analysis of the DNA binding (CG-1) domain of CAMTAs in plants and animals showed strong conservation of several amino acids. We mutated several conserved amino acids in the CG-1 domain to investigate their role in CAMTA3 function. Electrophoretic mobility shift assays using these mutants with a promoter of its target gene identified critical amino acid residues necessary for DNA binding activity. Furthermore, transient assays showed that these residues are essential for the CAMTA3 function in activating the *RSRE* (Rapid Stress Response Element)-driven reporter gene expression. In line with this, transgenic lines expressing the CG-1 mutants of CAMTA3 in the *camta3* mutant failed to rescue the mutant phenotype and restore the expression of CAMTA3 downstream target genes. Collectively, our results provide biochemical and genetic evidence that the transcriptional activity of CAMTA3 is indispensable for its function.

## Introduction

Plants in their environment are continuously subjected to biotic and abiotic stresses, which limit plant growth, and crop yield (Dhlamini et al., 2005; Miller et al., 2010). Therefore, plants have evolved various physiological and biochemical mechanisms for stress adaptation that rely mainly on changes in gene expression patterns (Fujita et al., 2006; Hadiarto and Tran, 2011). In this process, transcription factors (TFs) represent the master switches that target stress-responsive genes and regulate their expression (Brivanlou and Darnell, 2002). The Calmodulin-Binding Transcription Activators (CAMTAs) family is one of the well-characterized calmodulin (CaM)-binding TFs that work downstream of the calcium signaling pathway to elicit plant responses to various biotic and abiotic stresses (Reddy et al., 2000; Yang and Poovaiah, 2000). CAMTAs have been identified in plants, humans, Drosophila as well as worms and are evolutionarily conserved from plants to humans (Bouche et al., 2002; Yang and Poovaiah, 2002; Choi et al., 2005; Han et al., 2006; Song et al., 2006; Finkler et al., 2007). A typical CAMTA protein has various ordered functional domains with amino acid sequence stretches that are evolutionarily conserved (Finkler et al., 2007; Rahman et al., 2016; Iqbal et al., 2020). These domains include a DNA-binding domain (CG-1), a non-specific DNA-binding domain (TIG), ankyrin repeats (ANK) for protein-protein interaction, followed by five tandem repeats of a Ca^2+^-independent CaM-binding domain (IQ), and a Ca^2+^-dependent CaM-binding domain (CaMBD).

In Arabidopsis, there are six CAMTAs (CAMTA1 to CAMTA6), which are differentially expressed in response to multiple stresses (Reddy et al., 2000; Yang and Poovaiah, 2002). Among these CAMTAs, CAMTA3 (also known as signal-responsive1, SR1) is the most studied member and regulates diverse biotic and abiotic stress responses, and thigmomorphogenesis (Finkler et al., 2007; Du et al., 2009; Kim et al., 2017; Wang et al., 2021; Yuan et al., 2021; Darwish et al., 2022; Zeng et al., 2022)). Several transcriptomic studies comparing WT and *camta3* mutants revealed a dual regulatory role for CAMTA3 in multiple cellular pathways (Galon et al., 2008; Kim et al., 2013; Prasad et al., 2016). Mutants of *camta3* exhibit up-regulation of plant immune, SA biosynthetic, and salt stress-responsive genes (Galon et al., 2008; Du et al., 2009; Prasad et al., 2016) while genes associated with herbivory and low temperature stresses showed down-regulation (Doherty et al., 2009; Prasad et al., 2016). In line with this, *camta3* mutants exhibited autoimmune phenotypes including stunted growth, chlorosis and leaf lesions, accumulation of SA, constitutive expression of defense genes and enhanced resistance against bacterial and fungal pathogens (Galon et al., 2008; Du et al., 2009; Nie et al., 2012; Rahman et al., 2016). Our studies have shown that CAMTA3 functions as a negative regulator for salt tolerance in Arabidopsis (Prasad et al., 2016). On the other hand, CAMTA3 acts as a positive regulator for cold and drought stress response as well as defense against insect herbivory and wound-induced response (Laluk et al., 2012; Qiu et al., 2012; Zeng et al., 2022). CAMTA3 works in concert with CAMTA1, CAMTA2 and CAMTA5 to increase plant freezing tolerance by inducing the expression of the *CRT/DRE Binding Factor (CBF*) genes and other cold-induced genes (Doherty et al., 2009; Kim et al., 2013; Kidokoro et al., 2017). The *camta3* mutants were also more susceptible to herbivore attack and showed decreased content of the herbivory deterrents, glucosinolates as compared to the WT plants (Laluk *et al*., 2012). Moreover, CAMTA3 has been recently reported to be a positive regulator for the suppression of plant viral infection by activating the RNAi defense machinery (Wang et al., 2021). Therefore, CAMTA3 functions both as a transcriptional repressor for defense- and salt stress-related genes as well as a transcriptional activator for genes involved in cold, drought, glucosinolate metabolism, herbivory, wound-induced and general stress responses (Doherty et al., 2009; Laluk et al., 2012; Qiu et al., 2012; Benn et al., 2014; Prasad et al., 2016).

The CAMTA CG-1 domain binds to *CGCG* or *CGTG* core motifs with the *CGCG*-core consensus motif (*A*/*C*)*CGCG*(*C*/*G*/*T*), whereas the *CGTG*-core consensus motif is (*A*/*C*)*CGTGT* (Yang and Poovaiah, 2002; Galon et al., 2008; Doherty et al., 2009; Du et al., 2009; Galon et al., 2010). Furthermore, CAMTA3 has been shown to act as a transcriptional activator of early general stress response genes by binding to the *Rapid Stress Response Element* (Copin et al.) element that contains the *CGCGTT* motif (Walley et al., 2007; Benn et al., 2014). Moreover, it has been suggested that CAMTA3 regulates plant immunity by modulating the expression of the defense genes; *EDS1*, *NDR1*, *EIN*3 and *NPRI* by binding to the *CGCG* elements in their promoters (Du et al., 2009; Nie et al., 2012; Yuan et al., 2021) it is the thought that CAMTA3 regulates plant immunity by regulating the expression of defense genes. However, Lolle *et al*., (Lolle et al., 2017) have shown that the enhanced defense phenotype observed in *camta3* is primarily because of the activation of the NLRs (Nucleotide-binding domain Leucine-rich Repeat), DSC1 and DSC2, rather than the loss of the transcriptional activity of CAMTA3 as a negative regulator of gene expression. In a genetic screen, they identified two dominant-negative NLRs; *DSC1-DN* and *DSC2-DN* that suppressed the autoimmune phenotype of the *camta3* mutant (Lolle et al., 2017). Expression of *DSC1-DN* or *DSC2-DN* in *camta3* mutants suppressed the autoimmune phenotype, restored the expression of the defense genes; *PR1* and *EDS1* to WT levels and repressed plant resistance to fungal and bacterial pathogens (Lolle et al., 2017). Furthermore, expression of *DSC1* or *DSC2* in *Nicotiana benthamiana* triggered the HR cell death response, but co-expression of *CAMTA3* prevented this (Lolle et al., 2017). Furthermore, BiFC and FRET assays showed CAMTA3 and DSCs to be a part of a nuclear localized complex (Lolle et al., 2017). Together these results led to the suggestion that DSC1/DSC2 and CAMTA3 form a guard-guardee complex, similar to that observed with RPM1 and RPS2 NLR and R1N4 proteins (Jones et al., 2016). Based on these results, they suggested that modification of CAMTA3 by effectors or loss of CAMTA3 triggers the corresponding NLR guards to activate plant defense genes (Lolle et al., 2017). Here, we tested if the DNA binding and transcriptional activity of CAMTA3 is necessary for its function in plant immunity. We created point mutations in the conserved amino acids of the CAMTA3 CG-1 domain, evaluated the DNA binding and transcriptional activity of the mutated versions, and tested their ability to complement the *camta3* mutant phenotype. Our results show that the CAMTA3 DNA binding activity of the CG-1 and its modulation of transcription is essential for its function in plant immunity. Furthermore, our study identified amino acids in the CG-1 domain that are critical for its DNA binding activity.

## Materials and Methods

### Identification and selection of the conserved amino acids in the CG-1 domain for mutational studies

The genomic and full-length protein sequences of 1343 CAMTA transcription factors (including the splice variants) belonging to 165 plant species obtained from the “Plant Transcription Factor Database” (http://planttfdb.gao-lab.org) were used for analysis. A full-length protein sequence alignment across the various plant species was generated using the “T-Coffee” software as described at http://planttfdb.gao-lab.org. The protein sequences from multiple species were truncated to include only the CG-1 domain, which were then aligned using the Hidden Markov Model-Guided method (http://planttfdb.gao-lab.org) to generate a sequence logo showing the highly conserved amino acids (Jin et al., 2017). Furthermore, the Human HsCAMTA (HsCAMTA1, Q9Y6Y1) and the *Drosophila* dmCAMTA (ABI94369) amino acid sequences were obtained from the NCBI database, and their CG-1 domains were compared to that of the AtCAMTA3 sequence. Based on the results of these alignments, some of the amino acids are highly conserved across all plants, human and insects, and these were selected as candidates for mutational studies.

### Generation of CAMTA3 CG-1 domain mutants

Total RNA was isolated from Arabidopsis using the Plant RNAeasy Kit (Qiagen, USA). About 2 µg of DNAse-treated total RNA was used for the first strand cDNA synthesis using Superscript III (Invitrogen, USA) as per manufacturer’s instructions. The full-length *CAMTA3* cDNA was PCR amplified using the HS Primestar DNA polymerase (Takara, USA) with forward and reverse primers bearing *BamH*I and *Xho*I, respectively. The amplified product was gel extracted, digested with *BamH*I and *Xho*I enzymes and cloned in *pET28a* digested with the same enzymes. After verifying the sequence, the full-length coding sequence clone of *CAMTA3* in *pET28a* was used as a template for the generation of the mutants.

Mutants in the CG-1 domain were created using the Q5 site-directed mutagenesis kit (New England Biolabs, USA) as per manufacturer instructions. The primers used for creating the mutations were designed using the online design tool, NEB Base changer web portal (https://nebasechanger.neb.com). The primers were designed with 5’end annealing back-to-back, wherein the forward primer incorporated the mutated base(s) (Table S1). The generated plasmids carrying the mutations in the CG-1 domain were subsequently transformed into high-efficiency *E. coli* competent cells for amplification. The mutants were then sequenced using different primers to cover the entire length of the *CAMTA3* coding sequence.

### Expression and purification of the CG-1 domain in E.coli

#### Protein expression and purification

For expression of the CG-1 domain (153 aa) in *E. coli*, the *CAMTA3 CG-1* domain (1-459 bps) was PCR amplified from the *pET*-28a plasmid bearing WT or individual six-point mutations (M1 to M6) using HS Primestar DNA polymerase (Takara) with forward and reverse primers bearing *BamH*I and *Xho*I, respectively. The amplified product was gel extracted, digested with *BamH*I and *Xho*I enzymes and cloned in *pET28a* digested with the same enzymes. After verifying the sequence, the *pET*-28a plasmid bearing WT CG-1 domain or its six-point mutants (M1 to M6) were transformed into BL21-Codon Plus (DE3)-RIL (Stratagene, USA) cells. The single transformed colony was inoculated into 5 ml of LB medium with kanamycin (50 mg/L). The culture was then incubated at 37°C in a shaker maintained at 200 rpm for 15h. About, 500 µl of overnight culture was added to 200 ml of liquid LB medium supplemented with 50 mg/L of kanamycin and incubated at 37°C with vigorous shaking till an A_600_ of 0.5 to 0.6 was reached and then induced with 0.1 mM IPTG for 4 h at 37°C. The cells were harvested by centrifugation at 8000 rpm for 15 min at 4°C. The pelleted cells were quickly washed with lysis buffer prior to the preparation of the extracts. The pelleted cells were resuspended in 20 ml of lysis buffer (20 mM HEPES-KOH pH8.0, 1M NaCl, 2 mM ß-ME, one tablet of EDTA-free complete Protease Inhibitor Cocktail, 100 mM PMSF, 1mM imidazole). The resuspended cells were sonicated four times with a pulse interval of 10 sec followed by 2 min incubation on ice. The cell extract was clarified by centrifugation (11,000 rpm, at 4°C for 60 min). This clarified extract was used for the purification of the His-tagged proteins using Ni-NTA agarose resin (Qiagen, USA). The agarose resin was washed twice by resuspending in lysis buffer followed by centrifugation at 3000 rpm and removing the supernatant. The clarified cell lysate was incubated with washed Ni-NTA agarose resin on a rotatory shaker for 1h at 4°C. After incubation, proteins bound to the resin were collected by centrifugation at 4°C, 500xg for 5 min. The resin was repeatedly washed by suspending it in wash buffer 1 (lysis buffer +10 mM imidazole) followed by incubation on a rotatory shaker for 5 min. This step was repeated till the Abs_280_ of the supernatant reached zero. This was followed by a final wash with wash buffer II (Lysis buffer + 20 mM imidazole). For elution of the His-Tagged proteins, the washed resin was resuspended in wash buffer II and transferred to the column and allowed to settle down. His-tagged proteins were eluted using elution buffer (lysis buffer +250 mM imidazole). Depending upon the bead volume, multiple fractions of fixed volume were collected. His-tagged proteins in the fractions were detected by Bradford assay. The membrane containing the purified proteins was probed with an anti 6X His Tag monoclonal antibody (Abcam; ab49746) conjugated to alkaline phosphatase at 1:2000 dilution. The fractions that exhibited single bands of the expected size were pooled and dialyzed overnight at 4°C against dialysis buffer (20 mM HEPES-KOH pH 8.0, 200 mM NaCl, 20% glycerol, 1mM DTT) in a dialysis membrane bag (3500 kDa cut off) (Spectrum, USA). The dialyzed His-tagged proteins were aliquoted and flash frozen in liquid nitrogen and stored at -80°C.

The flash-frozen purified proteins were thawed on ice and the protein concentration was estimated using Bradford reagent (Bio-Rad, USA). Ten µg of the purified protein was boiled in 1X sample buffer, resolved on 12% polyacrylamide gel with 10% SDS and electro-blotted onto a PVDF (Bio-Rad, USA) membrane. The blot was blocked overnight with 5% nonfat milk in TBST buffer (50 mM Tris-HCl, pH 7.5, 150 mM NaCl, 0.05% Tween-20). The membrane was then probed with an anti 6X His Tag monoclonal antibody (Abcam; ab49746) conjugated to alkaline phosphatase at 1:2000 dilution and detected with an alkaline phosphatase detection system.

#### EMSA assays

Electrophoretic mobility shift assay (EMSA) for studying DNA protein interaction was performed with purified CG-1 (DNA binding domain) variants using a non-isotopic method. For this purpose, the LightShift Chemiluminescent EMSA kit (Thermo Fischer Scientific, USA) was used as per the manufacturer’s instructions. The DNA probe was made by combining two oligos with one being 3’end biotin labeled containing the binding site of CAMTA3. The binding studies were conducted by incubating the 3’end labeled DNA probe (20 fmol) with the purified protein in a final reaction volume of 20 µl consisting of 1X binding buffer, 50 ng/µl Poly (dI. dC) with or without 4pmol of unlabeled DNA at room temperature for 30 min. The reaction was terminated using 1X loading dye, was resolved on 5% native polyacrylamide gel (in 0.5X TBE) electrophoresis and electro-blotted onto a nylon membrane. Subsequently, the membrane was cross-linked at 120 mJ/cm^2^ using a UV cross-linker with 254 nm bulbs for 1min. The membrane was developed using a chemiluminescent detection module (Thermo Fischer Scientific, USA) according to the manufacturer’s instructions. Briefly, the membrane was blocked with blocking buffer for 15 min and transferred to a blocking solution supplemented with stabilized Streptavidin-Horseradish Peroxidase Conjugate and incubated on a shaker for an additional 15 min. Subsequently, the membrane was washed four times with 1X wash buffer and incubated in substrate buffer for 5 min. The chemiluminescent substrate solution was poured onto the membrane and incubated for 5 min in dark. The membrane was wrapped in saran wrap and exposed to a gel documentation system equipped with a CCD camera (BioRad, USA) to capture the signal.

To determine the minimum amount of protein required for the binding to the probe, EMSA assays were performed using different protein concentrations in the presence of a constant amount of the labeled DNA. The chemiluminescence signal for each band was quantified using gel documentation software (Image Lab, BioRad, USA), and the fraction bound was plotted as a function of protein concentration.

### Generation of effector and reporter constructs

#### Effector constructs

The WT as well as the mutated versions of *CAMTA3* CG-1 mutants (*M1*-*M6*) were PCR amplified using the primers indicated in Table S1. The fragments were then cloned into *pFGC5941* binary vector (downstream to *CaMV35S* promoter) between the *Asc1* and *BamH1* sites using the restriction sites for *Asc*1 and *BamH*1 that were added to the forward and reverse primers, respectively. All the generated constructs were sequence verified, then each construct was transformed into *Agrobacterium tumefaciens* strain GV3101 and subsequently used for the generation of transgenic lines and transient assay experiment.

#### Reporter constructs

For making the reporter constructs for the transient assay experiments, a previously reported *4X RSRE* sequence together with the upstream sequence of a minimal NOS promoter (-101 to +4) (Walley et al., 2007) (*4XRSRE NOS*) was synthesized. A mutated version (*4XmRSRE NOS*) was also synthesized by changing the core binding motifs (Suppl. Fig. 4). The *4XRSRE NOS* sequence was synthesized with *Pst*I and *Stu*I restriction sites at the 5’ end and *Sal*I and *Nco*I at the 3’ end to facilitate cloning upstream of the luciferase reporter gene. The synthesized DNA fragments were amplified using high-fidelity PrimeStar HS DNA polymerase with the end primers (Table S1) that contained the above-mentioned restriction sites. The amplicons were gel purified and digested with *Pst*I and *Sal*I, and then used for replacing the *CaMV35S* promoter upstream of the luciferase gene in the pUC18 vector, resulting in pUC18 *4XRSRE NOS::LUC* or *4XmRSRE NOS::LUC* expression constructs. The resulting constructs were then digested with the *Sac*I and the ends were blunted with the Fast End Repair Kit (Thermofisher, USA) followed by another digestion with *Stu*I to obtain blunted ends fragments consisting of *4XRSRE NOS::LUC* or *4XmRSRE NOS::LUC*. These fragments were then inserted into binary vector *pFGC5941*, which was previously digested with *Stu*I and *Sma*I. The resulting plasmids were *pFGC5941 4XRSRE NOS::LUC-OCS* and *pFGC5941 4XmRSRE NOS::LUC-OCS*.

### Generation of the Transgenic lines

Each of the generated *pFGC5941* constructs with WT or CG-1 mutants of CAMTA3 was transformed into *Agrobacterium tumefaciens* and subsequently used to transform *camta3* mutant plants using the floral dipping method. The transgenic plants were then selected on MS plates containing Basta (10 µg ml^-1^) and genotyped by RT-PCR using the primers listed in Table S1. The selected plants were then selfed to obtain homozygous lines and three independent lines were chosen for each construct.

#### RNA Extraction and Gene Expression Analysis

For RNA isolation, leaf tissue was collected from 4-week-old plants grown in soil and flash-frozen in liquid nitrogen. Total RNA was isolated using TRIzol reagent (Invitrogen, USA) and treated with an RNase-free DNase (Promega, USA) to remove any genomic DNA contamination. Two µg of the DNAse-treated RNA was reverse transcribed into cDNA using oligo dT primer and Superscript II reverse transcriptase (Invitrogen, USA) according to the manufacturer’s instructions. The cDNA was diluted with 80 µl sterile nuclease-free water. Expression analysis was performed using RT-qPCR in a Roche LC480 machine (Roche, USA) using the preprogrammed “SYBR green-I 96 well program”. For every qPCR reaction, 10 µl of 2X LightCycler 480 SYBR Green I Master mix (Roche, USA) was used along with 1 µl of 5 µM of each primer and 2.5 µl cDNA template in a final reaction volume of 20 µl. *ACTIN2* was used as a reference gene as this gene does not exhibit any difference in its expression level among the various genotypes under different conditions. Fold change in expression was calculated and plotted with respect to control treatments. Three biological replicates were used for each experiment. Primers (see Table S1) for Real-time qPCR (RT-qPCR) were designed using the Primer Quest web tool (http://www.idtdna.com/Primerquest/Home/Index) from IDT (USA).

#### Transient Expression Assay in Nicotiana benthamiana

For the transient expression assay, we used *Agrobacterium tumefaciens* cells containing *CaMV35S::CAMTA3-OCS or CaMV 35S::CAMTA3-OCS* mutants (*M1-M6*) as effector constructs and *pFGC5941 4XRSRE NOS::LUC:OCS or pFGC5941 4XmRSRE NOS::LUC-OCS* as reporter constructs. The *Agrobacterium* cultures were grown for 36 h to OD600 ∼1.0-1.2. Individual cultures were then centrifuged at 3000 rpm for 20 min. Supernatants were then removed and the pellets were re-suspended in 10 ml infiltration medium (2mM Na_3_PO_4_, 50 mM MES, 0.5% glucose, 100 μM Acetosyringone) followed by incubation at room temperature for 3 h. Each culture was then adjusted to OD600=1.0 and the transgenes to be co-expressed were mixed in a 1:1 ratio. Each combination was then mixed with P19 culture in a 1:1 ratio before infiltration to suppress post-transcriptional gene silencing (PTGS) and enhance transient expression (Voinnet et al., 2003). Specifically, the *RSRENOS::LUC* construct or its dysfunctional mutant versions, *mRSRENOS*::*LUC*, were infiltrated independently in *Nicotiana benthamiana* leaves, alone or together with *CaMV35S::CAMTA3* or one of its mutant versions (M1-M6). The combined cultures were spot-infiltrated into six-weeks-old *Nicotiana benthamiana* leaves. Three plants (with three leaves per plant) were infiltrated with each combination. After 3 days, infiltrated leaves were collected, and flash-frozen in liquid nitrogen for protein extraction and quantification of LUC activity.

#### Protein Extraction and Quantification of LUC Activity

Infiltrated leaves were flash-frozen and ground to a fine powder in Tissue-Lyser and suspended at room temperature in 1X CCLR (25mM trisphosphate (pH 7.8), 2mM DTT, 2mM 1,2-diaminocyclohexane-N,N,Ń,Ń-tetraacetic acid, 10% glycerol, 1% Triton® X-100) (Promega, USA) with further homogenization on a rocker. After cell lysis, the extract was clarified by centrifugation at 4°C for 10 min at 13,500 rpm. using standard assay conditions. The LUC activity was assayed in the supernatant using the Luciferase Assay System (Promega, USA). Briefly, 20μl of cell lysate was added to 100μl of the Luciferase Assay Reagent into a tube and vortexed for 10sec prior to placing it in a cuvette holder. The generated luminiscence was measured using a luminometer (Turner Designs, USA) programmed to perform a 2-second measurement delay followed by a 10-second measurement read. Protein concentration was determined using the Bradford reagent (Bio-Rad, USA).

#### Protein Extraction and immunoblot analysis

For extraction of total protein, the leaves of the 30-day-old plants that were flash frozen were ground to final powder in TissueLyser and dissolved in 500 µl protein extraction buffer (40 mM K2HPO4, 10 mM KH2PO4, 1 mg ml-1 ascorbate, 0.05% β-mercaptoethanol (v/v) 0.1% TritonX-100, 1 mM PMSF) containing 1% protease inhibitor cocktail (Sigma-Aldrich, USA). The extract was centrifuged for 10 min at 13,500 rpm at 4oC. Protein concentration was determined using the Bradford reagent (Bio-Rad). Forty µg of total protein from each sample was resolved in 12% SDS gel and blotted onto a PVDF membrane (Millipore, USA). The blot was blocked with 5% non-fat milk in TBST buffer (50 mM Tris–HCl, pH 7.5, 150 mM NaCl, 0.05% Tween-20). Then, the membrane was probed with an anti-CAMTA3 polyclonal antibody at a 1:500 dilution and detected with an antirabbit secondary antibody conjugated with horseradish peroxidase using the chemiluminescence detection system (Pierce, ThermoFischer, USA).

## Results

### Identification of highly conserved amino acids in the DNA binding

Arabidopsis CAMTA3, like other typical CAMTAs, also has five distinct domains, which include a DNA binding domain (CG-1), a TIG domain involved in non-specific DNA interaction and also involved in the protein dimerization, ankyrin (ANK) repeats implicated in protein-protein interaction, a Ca^2+^-independent CaM-binding domain (IQ domain), and a CaMB domain, a Ca^2+^-dependent CaM-binding domain (CaMB) (Fig.1a). Previous studies have shown that the CAMTA3 CG-1 domain directly interacts with the promoters of some CAMTA3-regulated genes. To identify conserved amino acids in the CG-1 domain and their function in DNA binding activity, we aligned the amino acid sequence of this domain of 1343 CAMTAs belonging to 165 species representing different taxonomic plant groups (Jin et al., 2017). The analysis revealed very high conservation of several amino acids with a bit score of over 4 (Fig. 1b & Suppl. Fig. 1). We also compared the amino acid sequences of the CG-1 domain of CAMTAs from human and fruit fly with that of Arabidopsis CAMTA3 and identified the highly conserved amino acid stretches (indicated as boxes) across the entire domain between plants and animals (Fig. 1c). Since these amino acids are evolutionarily conserved among phylogenetically diverse organisms, we postulated that these amino acids could be critical for CAMTA3 function and would be good candidates for point mutation to understand their role in the DNA-binding activity. Therefore, to generate point mutations, we have selected six amino acids across the entire stretch of the CG-1 domain that exhibit a high bit score of >4 and are also conserved among plants, human and *Drosophila* (Fig. 1d). These mutations are represented as M1 to M6 and the conserved amino acids replaced with alanine are shown in parenthesis next to each mutant: M1 (W26A), M2 (W75A), M3 (H88A), M4 (YY102/103/AA), M5 (W118A) and M6 (Y132A) (Fig.1d).

**Figure 1.**
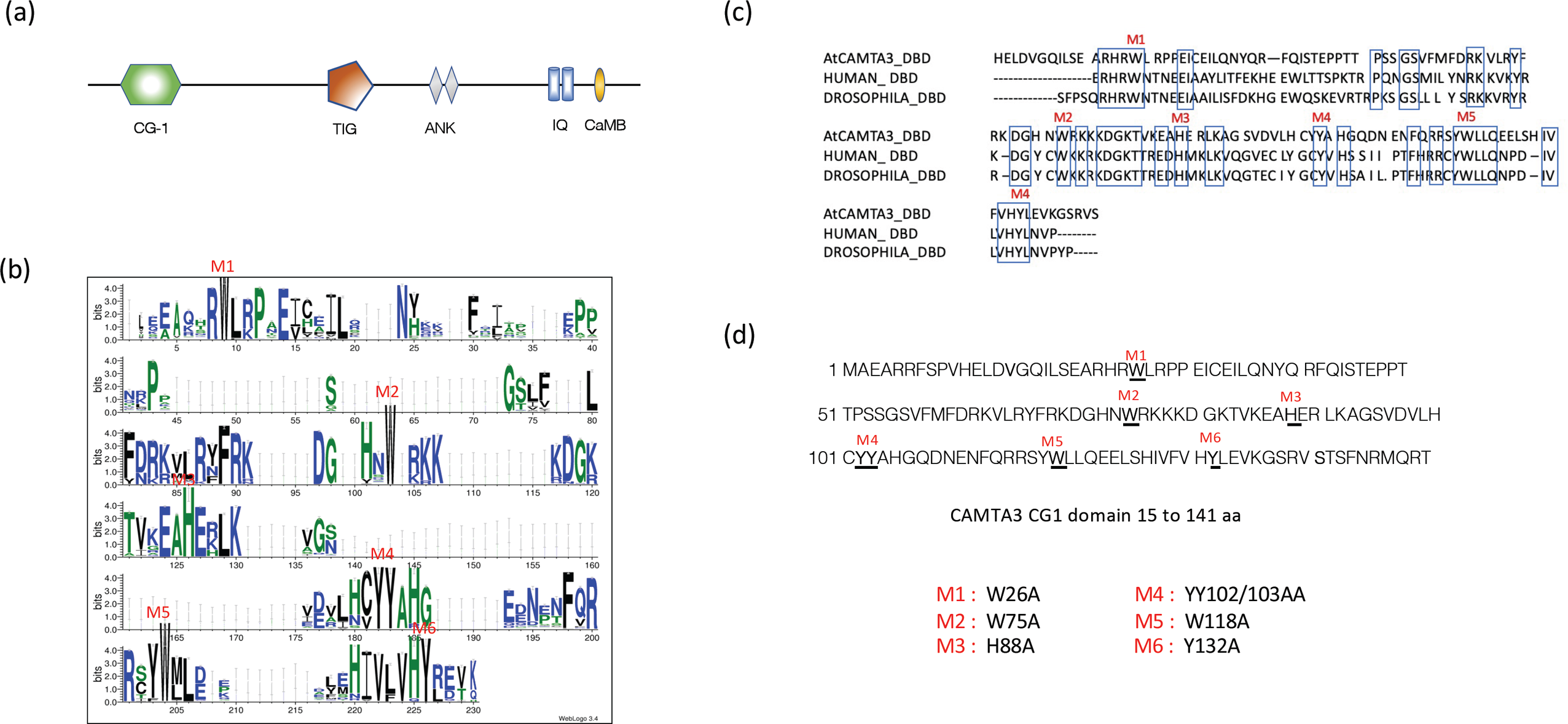
Identification of the evolutionarily conserved amino acid residues in the CG-1 domain of CAMTAs. **(a)** Schematic representation of full-length AtCAMTA3 protein indicating various domains. **(b)** Sequence logo of highly conserved amino acid residues in the CG-1 (DNA binding domain) of 1343 CAMTAs belonging to 165 plant species created. The web logo was generated using the Hidden Markov Model-guided method (Jin et al., 2017). **(c)** Alignment of amino acid sequence in the CG-1 domain of AtCAMTA3, HsCAMTA1 and DmCAMTA. Highly conserved amino acid residues are marked by blue boxes. **(d)** The amino acid sequence of the AtCAMTA3 CG-1 domain with the highly conserved amino acids (>4 bit scored) underlined. The letters M1-M6 indicated in (b-d) represent the conserved amino acids that are mutated to alanine (A).

### Identification of critical amino acids necessary for DNA binding activity of the CG-1 domain

To test the DNA binding activity of the CG-1 mutants, we performed EMSA assays using the CG-1 mutant proteins expressed in *E. coli* and a well-known CAMTA3 target DNA probe (shown in Fig. 2b). The wild-type and mutant proteins expressed in *E. coli* were purified using a His-tag column and their purity and size were confirmed through immunoblot analysis using an anti-His-tag antibody (Fig.2A). The immunoblot analysis showed a single band of an expected size (∼24 kDa) for all mutants (Suppl. Fig. 2). As CAMTA3 is known to play an important role in plant defense response, we chose the promoter of *PDF1.4* (a defense-related gene) as a candidate for designing the probe as it contains a well-characterized CAMTA3 CG-box binding site with “*CGCG”* core element (Yuan et al., 2018). Using electrophoretic mobility shift assay (EMSA), we initially performed binding studies with WT CG-1 recombinant protein in a concentration-dependent manner to confirm that the purified protein has DNA binding activity and determine the minimal protein concentration required for binding. As expected, EMSA showed a significant binding of the purified WT CG-1 protein to the *PDF1.4* probe in a concentration-dependent manner, indicated by a shift in the probe due to DNA-protein complex formation (Fig. 2b). Notably, a signal of DNA-protein complex was detected even at 25 ng protein. Our study indicated that a 50 ng or more protein was required for good binding to the *PDF1.4* probe *in vitro*. Based on these results, we chose a protein concentration of 500 ng for the EMSA experiment so that we can detect weak binding of CG-1 mutants to the probe. To further evaluate the binding and specificity of WT and CG-1 mutants (M1 to M6) to the *PDF1.4* probe, 10X unlabeled oligo of *PDF1.4* was used as a competitor in a separate reaction. Apart from WT and M5 mutant, none of the other CG-1 mutant proteins exhibited DNA binding (Fig. 3a), indicating that these mutated amino acids in the CG-1 domain are required for its binding to the target DNA. The inclusion of competitor oligo in the reaction significantly reduced the binding of the WT and M5 mutant to the target DNA, indicating highly specific interaction (Fig. 3a). However, despite the ‘W118’ amino acid is also highly conserved across the kingdoms, its mutation to ‘A’ in the M5 mutant showed DNA binding activity. Therefore, to rule out the possibility that binding of the M5 mutant to the probe was attributed to high protein concentration and to compare M5 activity with that of WT CG-1, we performed an EMSA using serial concentrations of M5 protein. As shown in Figure 3b and Supplementary Figure 3, the DNA-protein complex signal for the M5 mutant was detectable beginning with 100 ng protein concentration, which is four times more than the WT CG-1 protein concentration (25 ng). Furthermore, a significantly lower signal intensity level was observed with the M5 protein as compared to that observed with the WT protein, indicating a relatively weak binding for the M5 mutant with the target DNA.

**Figure 2.**
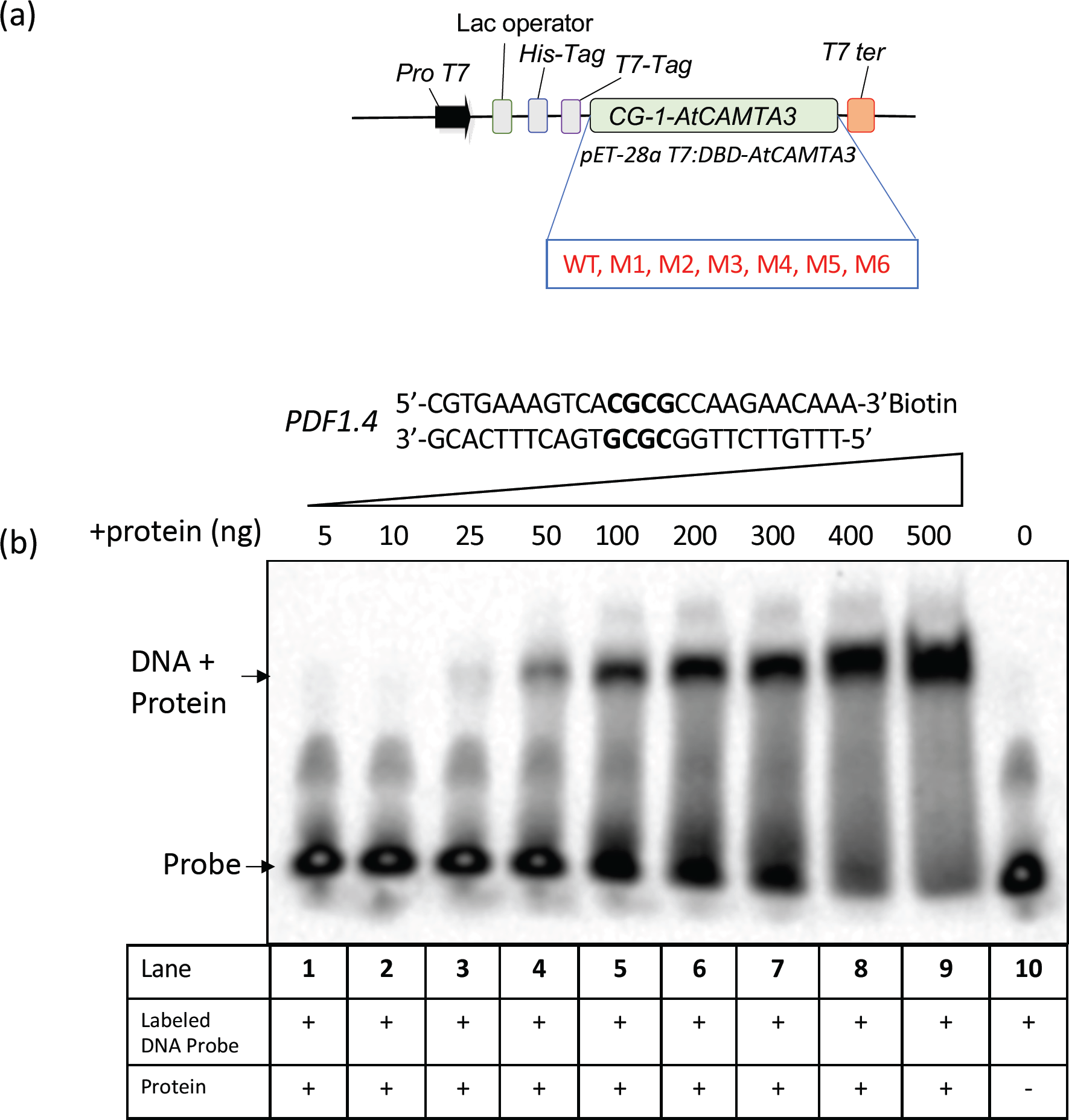
Electrophoretic mobility shift assay (EMSA) to determine the binding activity of WT CG-1 domain. **(a)** Schematic representation of the construct that was used for expression and purification of the WT and mutated CG-1 (DNA binding) domain variants in *E. coli*. The protein was purified using Ni-NTA agarose column. **(b)** Increasing concentrations of the purified recombinant WT CAMTA3 CG-1 protein (Lanes 1 to 9) were used for EMSA assays along with the *PDF1.4* probe to determine the minimal protein concentration required for binding. Probe alone is shown in lane 10.

**Figure 3.**
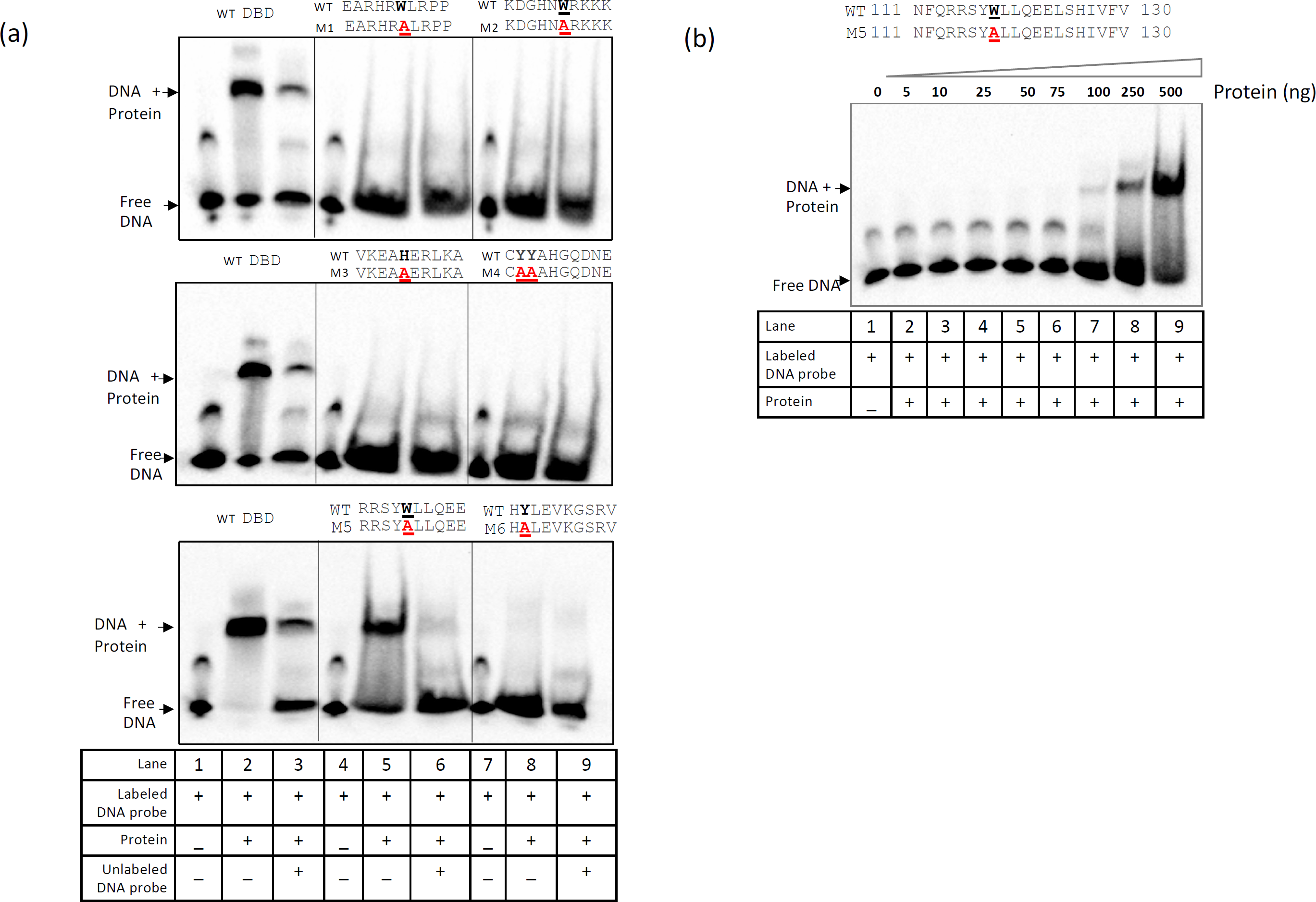
The majority of CG-1 point mutations abolished binding to the *PDF1.4* probe. **(a)** EMSA assays with purified WT and CG-1 mutant proteins using the *PDF1.4* probe. The majority of the CG-1 mutants (M1, M2, M3, M4 & M6) failed to bind to the *PDF1.4* probe as indicated by the absence of mobility shift of the labeled probe. Only the WT and the CG-1 domain variant (M5) with W118A mutation exhibited binding to the *PDF1.4* probe. The inclusion of competitor significantly reduced WT and M5 binding to the probe. **(b)** A protein concentration-dependent binding of M5 protein to *PDF1.4* probe. Note: the binding of M5 protein to the probe occurs only at concentrations of at or above 100 ng.

### DNA binding activity is required for transcriptional activation of stress-responsive reporter gene expression

Previous genetic screens have shown that CAMTA3 is an important regulator of rapid stress-responsive genes (Walley et al., 2007; Benn et al., 2014). It has been shown that CAMTA3 can activate the reporter gene (Luciferase; *LUC*) driven by the Rapid Stress Response Element (Copin et al.)-driven luciferase reporter (*RSRE::LUC* reporter) (Benn et al., 2014). Consistently, the *camta3* mutant exhibited reduced RSRE::LUC activity (Bjornson et al., 2014). Therefore, to further validate our in *vitro* binding results of CG-1 mutant, we transiently co-expressed *RSRE::LUC* construct or its mutated dysfunctional *RSRE* (*mRSRE*::*LUC*) (Fig. 4a) together with constitutively expressed *CAMTA3* (*CaMV35S::CAMTA3)* or individual *mCAMTA3* variants (*M1*, *M2*, *M3*, *M4*, *M4*, *M5* or *M6*) effector constructs (Fig. 4b) in *Nicotiana benthamiana* leaves. Consistent with our EMSA results, among all the CG-1 mutants, only the *M5* mutant behaved like WT *CAMTA3* and clearly displayed strong reporter gene activation. However, all other mutants (*M1*, *M2*, *M3*, *M4* & *M6*) exhibited highly reduced RSRE::LUC activity (Fig. 4c). Interestingly, in agreement with previous reports, reporter gene activation was only detected in leaves co-infiltrated with the *RSRE::LUC*, and no activation was detected in the assays with the mutated dysfunctional *mRSRE*::*LUC* construct. Taken together, these data imply that except for the *M5* mutant, the amino acid residues mutated in all the mutant lines (*M1*, *M2*, *M3*, *M4* & *M6*) are essential for the function of CAMTA3 in activating the *RSRE-*driven reporter.

**Figure 4.**
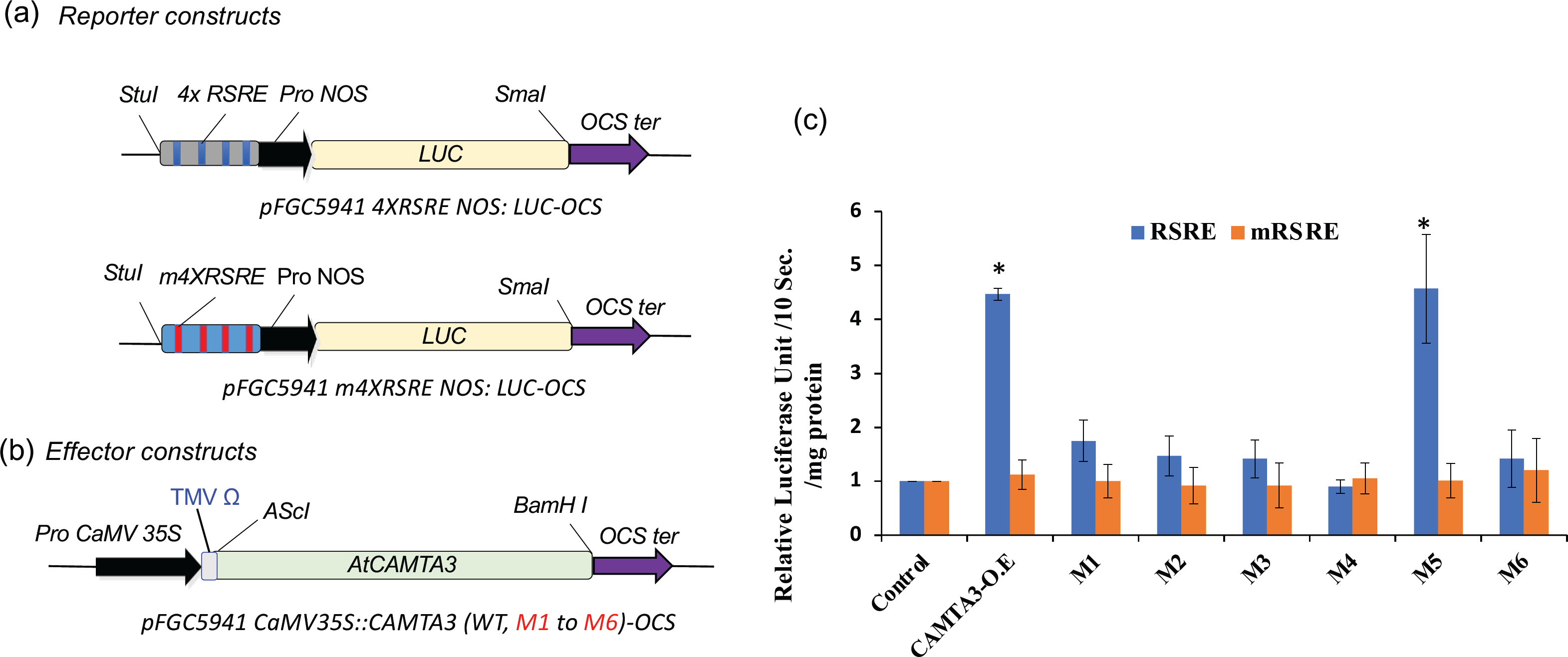
Full-length AtCAMTA3 CG-1 mutants curtailed the induction of *RSRE::LUC* reporter expression. Schematic representation of the reporter **(a)** and effector **(b)** constructs used in transient expression assays with *N. benthamiana* leaves for quantifying the induction of *RSRE::LUC* reporter expression by full-length WT and mutated AtCAMTA3. Reporter constructs include 4X *RSRE* elements (upper panel) and 4X *mRSRE* (lower panel). The effector constructs for expression of either full-length WT or CG-1 mutants (M1 to M6) of CAMTA3 driven by CaMV35S promoter are shown. **(c)** Transient expression of *RSRE::LUC* construct or its mutated dysfunctional *RSRE* (*mRSRE* ::*LUC)* alone (control) or together with *CaMV35S::CAMTA3 (CAMTA3-O.E)* or *mCAMTA3* variant (*M1*, *M2*, *M3*, *M4*, *M4*, *M5* or *M6*) in *Nicotiana benthamiana* leaves. Only the CAMTA3 WT and M5 mutant displayed strong *RSRE::LUC* activity. Whereas the CAMTA3 CG-1 mutant variants (M1, M2, M3, M4 & M6) exhibit curtailed induction of *RSRE::LUC* activity. Lines on top of the bars represent the standard error of the mean and asterisks above the line represent statistically significant differences (P<0.05) in reporter gene expression. The infiltration experiments were carried out three times for each construct with three independent plants.

### DNA binding activity is essential for CAMTA3 function in plant immunity

At 19–21°C, *Atcamta3* loss of function mutants exhibit autoimmune phenotype including reduced growth, chlorosis associated with leaf lesions and constitutive expression of the defense genes (Du et al., 2009). To investigate the impact of the CAMTA3 CG-1 domain mutations on the phenotype as well as the expression of CAMTA3 downstream target genes, we complemented the *camta3* mutant with WT or individual CAMTA3 CG-1 mutants (M1, M2, M3, M4, M5 or M6) (Fig. 5a). Immunoblot analysis of the protein from WT, CAMT3-YFP (CAMTA3-O.E.), *camta3* and lines complemented with the *CAMTA3* CG-1 mutants (M1 to M6) with an antibody specific to CaMBD of CAMTA3 indicated expression of the CAMTA3 in all lines except for *camta3* mutant (Fig. 5b). However, except for the mutant line *M5*, all mutant lines (*M1*, *M2*, *M3*, *M4* & *M6*) showed autoimmune phenotype similar to the *camta3* mutant phenotype (Fig. 5c) indicating that these mutant proteins are non-functional *in vivo*. Whereas the mutant line M5 showed a similar phenotype as that of the *CAMTA3* overexpression line.

**Figure 5.**
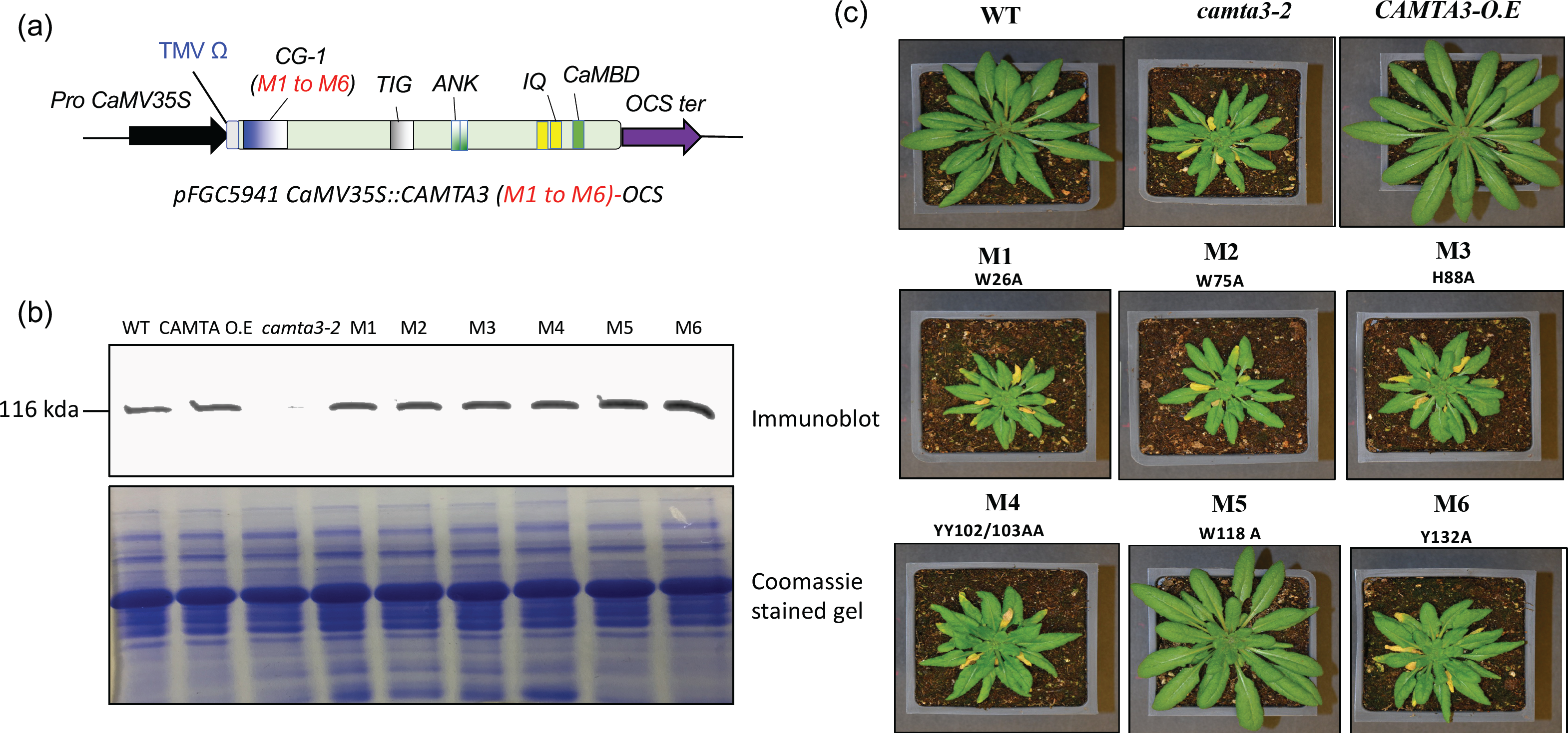
Expression and phenotypic analysis of CAMTA3 CG-1 mutants in *camta3* background. **(a)** Schematic representation of the constructs used for transformation of *camta3-2* (T-DNA insertion mutant) for expression of the WT and CG-1 mutant proteins of CAMTA3. **(b)** CAMTA3 protein levels in four-week-old WT and transgenic lines complemented with WT or CG-1 mutants (M1 to M6) of CAMTA3. The *camta3-2* plants transformed with *CaMV35S::CAMTA3 (CAMTA3-O.E)* and CG-1 mutants (M1 to M6) were grown under day neutral conditions at 19°C, and protein expression was detected using an antibody against CaMBD peptide. **(c)** Photographs of four-week-old plants depicting phenotypes of the WT, *camta3-2,* CAMTA3-O.E. and *camta3-2* plants expressing the mutants (M1 to M6).

Furthermore, in these lines, we analyzed the expression of some of the CAMTA3 downstream target genes that are differentially regulated by CAMAT3 (Prasad et al., 2016). We selected a few genes that are implicated in plant immunity (*CBP60, EDS1, ICS1 and ALPHA-DOX1*) and cold tolerance (*COR15* and *KIN1)* and analyzed their expression in CAMTA3 CG-1 mutant lines. It was shown previously that these defense response genes are upregulated whereas the cold response genes are downregulated in the *camta3* knockout mutant (Prasad et al., 2016) (Fig. 6). Similar to the *camta3* knockout mutant, the lines expressing *M1*, *M2*, *M3*, *M4* and *M6* showed enhanced expression of *CBP60, EDS1, ICS1 and ALPHA-DOX1* genes and suppressed expression of *KIN1* and *COR15* as compared to the WT. Whereas the mutant line *M5* showed expression patterns for these genes like the WT and *CAMTA3* overexpression lines. Taken together, these data imply that, except for the *M5* mutant, the amino acid residues mutated in all the mutant lines (*M1*, *M2*, *M3*, *M4* & *M6*) and the DNA binding activity are essential for the function of CAMTA3.

**Figure 6.**
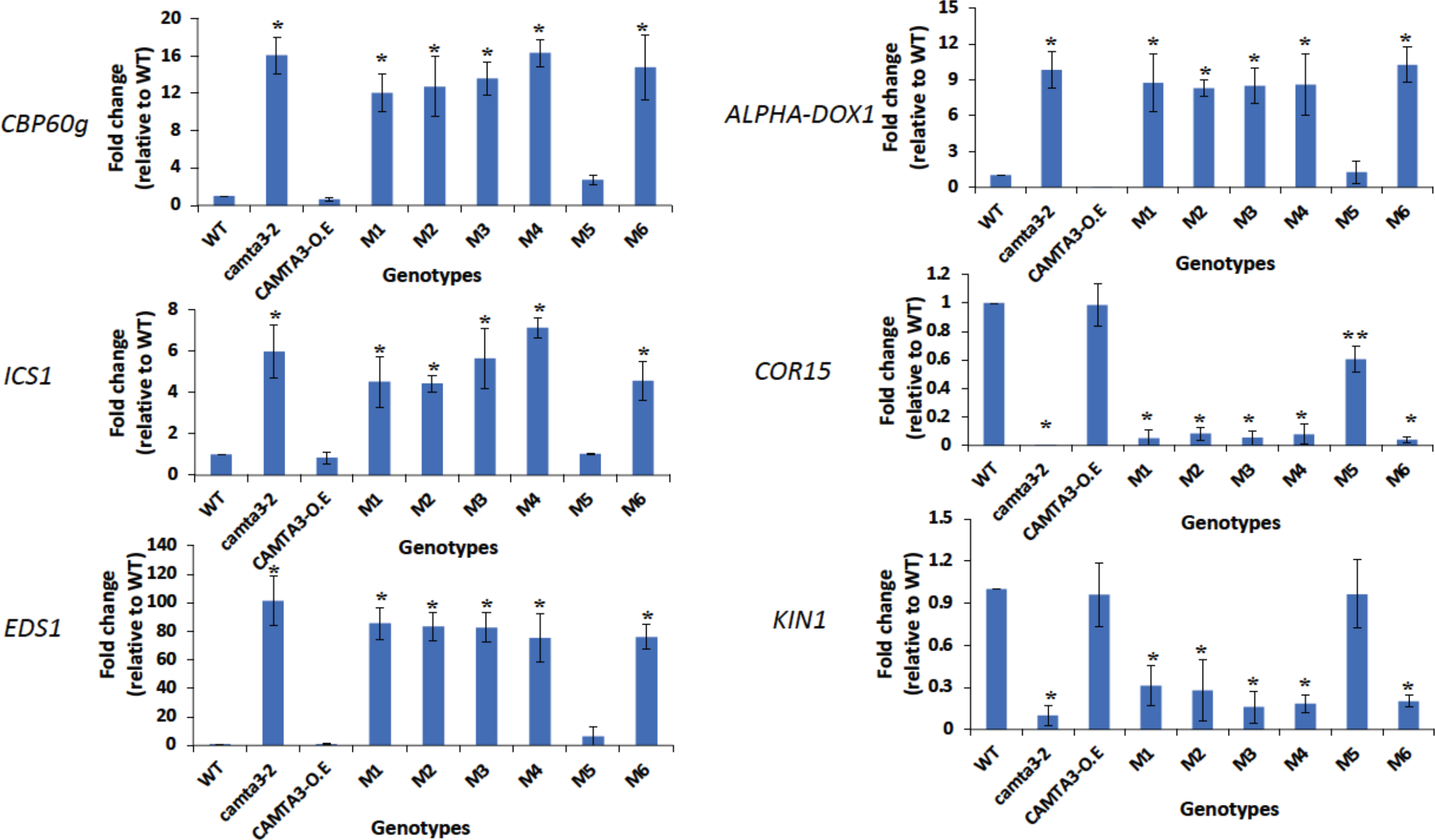
Repression of defense-related genes and activation of cold-responsive genes in *camta3-2* plants expressing the CAMTA3 CG-1 mutants. Comparison of transcript levels of the genes associated with plant defense (*CBP60g*, *ICS1*, *EDS1 ALPHA-DOX1*) and cold (*COR15* and *KIN1*) responses in WT plants and *camta3-2* mutants as well as *camta3-2* mutants expressing *CaMV35S::CAMTA3* (*CAMTA3-OE*) and *CG-1* mutants (M1 to M6). The plants were grown in a growth chamber at 20°C under 12:12 h light/dark cycles, with a light intensity of 200 μmol m^−2^ s^−1^and relative humidity of 50%. Leaves of six-week-old plants were harvested for RNA isolation. Statistically significant differences (P<0.05) in reporter gene expression in all lines compared to WT are shown with asterisks.

## Discussion

Several genetic and biochemical studies have shown that CAMTAs transcriptional regulatory activity is associated with their binding to the (*A/C/G*) *CGCG*(*T/C/G*), (*A/C*)*CGTGT* or *CG*(*C/T*)*G cis*-elements in the promoter regions of their target genes, primarily through the CG-1 domain (Yang and Poovaiah, 2002; Mitsuda et al., 2003; Choi et al., 2005; Finkler et al., 2007; Du et al., 2009). However, more recently, Lolle *et al*., (Lolle et al., 2017) suggested that CAMTA3 acts as a guardee for NLR immune receptors in modulating plant immunity rather than having a direct transcriptional activity role in regulating plant immunity. They have identified two TIR-type NLR proteins, DSC1 and DSC2 to be guards for the host effector target (guardee) CAMTA3. They suggested that the autoimmunity in *camta3* plants results from ectopic activation of the two NLRs: DSC1 and DSC2 rather than loss of the CAMTA3 transcriptional repressor function per se (Lolle et al., 2017). *camta1* or *camta2* null alleles in *camta3* mutants enhance the latter’s autoimmune phenotype, whereas neither *camta1* nor *camta2* shows an autoimmune phenotype (Kim et al., 2013). Interestingly the expression of *DSC1-DN* or *DSC2-DN* versions in the *camta1 camta3* double mutant didn’t rescue the autoimmune phenotype (Lolle et al., 2017). Therefore, if CAMTA3 works only as a guardee for DSC1, expression of the *DSC1-DN* or *DSC2-DN* into the *camta1 camta3* background, should be able to repress the autoimmune phenotype. In another study, the expression of a truncated version of CAMTA3^1–344^ that includes the CAMTA3 CG-1 domain suppressed the autoimmune phenotype of the *camta2 camta3* (Kim et al., 2017). Moreover, the expression of the *DSC1* orthologue of *Gossypium hirsutum* (*GhDSC1*) as well as *CAMTA3* was found to be induced in response to the *Verticillium* wilt and JA treatment in Arabidopsis, suggesting a coordinated *DSC1* and *CAMTA3* response in Arabidopsis to Verticillium wilt (Li et al., 2019). CAMTA3 has also been reported as a negative regulator for plant immunity by direct binding to the promoter regions of the SA positive regulator *CBP60g* as well as the TNL- and CNL-mediated defense pathways regulators, *EDS1* and *NDR1*, respectively and suppressing their expression (Du et al., 2009; Nie et al., 2012; Rahman et al., 2016). Additionally, Doherty *et al*., (Doherty et al., 2009) demonstrated that the CG-1 domains of CAMTA1, 2, 3, and 5 bind to the *CM2* motif in the promoter region of *CBF2*. They also provided evidence that CAMTA3 acts as a positive regulator for the cold-induced *CBFs* gene expression in Arabidopsis (Kidokoro et al., 2017).

One way to address whether DNA binding and transcriptional activities of CAMTA3 are necessary for regulating plant immunity is to generate CAMTA3 mutants that are unable to exhibit DNA binding and thus abolish their transcriptional activity. Irrespective of the plant species, virtually all CAMTAs have a CG-1 domain in the N-terminal region of the protein, indicating that the position of this domain is evolutionarily conserved (Bouche et al., 2002; Finkler et al., 2007; Rahman et al., 2016). The CG-1 domain is also present in human and animals CAMTAs (Finkler et al., 2007) (Fig. 1). Our analysis of the CG-1 domain revealed a high conservation of aromatic amino acids such as W, H, Y and F in the CG-1 domain indicating their likely role in DNA binding (Baker and Grant, 2007). Similarly, a high occurrence of these W and Y amino acids was also noted in the DBD of WRKY, ERF, MYB and MYB-related TFs that are also involved in plant immune response (Shoji et al., 2013). Since *camta3* mutants exhibit temperature-dependent constitutive autoimmune response, we hypothesized that the necessity of CAMTA3 transcriptional activity can be effectively scored by expressing WT and CAMTA3 CG-1 mutant variants in *camta3* background and see if it will rescue the mutant phenotype. Remarkably, the majority of the CG-1 mutant variants (M1-M4 and M6) that exhibited no DNA binding activity also failed to rescue the mutant phenotype, while WT and M5 with DNA binding activity did complement the phenotype (Fig.5). Together, these results suggest that the transcriptional activity of CAMTA3 is required for its function in plant immunity. We further validated this by analyzing the expression of several CAMTA3-regulated genes that were previously reported to be involved in plant immunity and cold response (Doherty et al., 2009; Du et al., 2009; Nie et al., 2012; Prasad et al., 2016; Kim et al., 2017). The expression levels of all the immune and cold response genes in the M1 to M4 and M6 variants were similar to that in the *camta3* mutant, whereas their expression in the M5 mutant was like WT plants (Fig.6). Furthermore, to exclude the possibility that some of these results were due to the *in vivo* destabilization of CAMTA3 proteins as a result of the mutation, we analyzed the proteins levels and compared them with that of WT. Our results revealed similar levels of protein in all the CG-1 mutants and WT plants, implying that these CG-1 mutations did not alter the stability of the CAMTA3 protein. Collectively, our results provide strong biochemical and genetic evidence that the transcriptional activity of CAMTA3 is essential for its function. The requirement of transcriptional activity of CAMTA3 for its repressor and activator function was further supported by the function exhibited by the NRM module (CAMTA^1-334^), a truncated version of CAMTA3 protein that includes the N terminal CG-1 DNA binding domain and lacks other domains (Kim et al., 2017; Chao et al., 2022). Expression of NRM in *camta2 camta3* double mutant not only suppressed the expression of *ICS1*, *CBP60g*, *PRI* but also rescued the autoimmune phenotype of this mutant (Kim et al., 2017). In another study, NRM (CAMTA3^334^) was also reported to be sufficient to rapidly induce the expression of *CBF2* and two other cold-inducible genes; *EXPansin-Like A1* (*EXPL1*) and *Nine-Cis-Epoxycarotenoid Dioxygenase 3* (*NCED3*) under cold stress (Kim et al., 2017; Chao et al., 2022). However, these studies did not identify the amino acids involved in the DNA binding activity of the NRM. In this context, our study provides evidence for the role of the amino acids involved in the transcriptional activity of CAMTA3.

As CAMTA3 stability and turnover are known to be critical for its function as a repressor for the immune response (Zhang et al., 2014; Jiang et al., 2020), we show that the minimum concentration of 25 ng of WT CAMTA3 CG-1 domain to be required for binding to *PDF1.4* (Fig. 2). However, most of the CAMTA3 CG-1 mutants (M1, M2, M3, M4 and M6) failed to bind to the same probe even at a higher protein concentration of 500 ng (Fig.3) indicating that these amino acids are critical for DNA binding and CAMTA3 transcriptional activity. Interestingly, the M5 mutant exhibited binding to the probe, albeit at higher concentrations (100 ng), indicating weak binding (Fig.3). Previously, it has been shown that the expression of the truncated variant of CAMTA3 comprising the CG-1 domain (334 amino acids, NRM), exhibited transcriptional activation and repressor activities *in planta* (Kim et al., 2017). Interestingly, mutations of K108A and K141E in the CG-1 domain of the human CAMTA2 and CAMTA1, respectively (Fig. 1), completely abolished their DNA binding activity indicating a critical role for these amino acids in the DNA-binding activity of these CAMTAs (Long et al., 2014). Indeed, this amino acid (K) is also conserved across the plant kingdom, however, the degree of conservation is slightly lower than that observed with other aromatic amino acids (Fig.1). However, the role of K in DNA binding cannot be ruled out in plant CAMTAs, and additional mutational studies are needed.

Previously, CAMTA3 has been shown to bind to the *RSRE* elements and activate downstream genes (Benn et al., 2014). Expression of full-length WT and CAMTA3 mutant proteins transiently in tobacco and tested their ability to activate the expression of luciferase gene driven by *RSRE* (*vCGCGb*). In agreement with earlier reports, the binding of WT CAMTA3 is highly specific, and it transcriptionally activates the luciferase gene expression. However, except for the M5 mutant, all other mutants failed to activate the luciferase reporter gene expression (Fig.4) which concurred with our results in figures 2 and 3, with truncated CAMTA3 proteins.

In conclusion, our studies with multiple mutants in the CG-1 domain of CAMTA3 indicate that the DNA binding and transcriptional activities of this protein are essential for its function in plant immunity. The current evidence suggests that CAMTA3 performs a dual role as a guradee of NLRs (DSC1/2) and as a transcriptional regulator. Further studies are needed to elucidate the mechanistic details of this dual role of CAMTA3. It is possible that elicitor-mediated modification of CAMTA3 may not only activate the NLR-mediated pathway but also regulate the transcriptional activity of CAMTA3.

## Acknowledgements

This work was supported by a grant from the National Science Foundation (MCB # 5333470) to ASNR

## Author contributions

A.S.N.R. directed the project. K.V.S.K.P., A.A.H., and A.S.N.R. designed the experiments. K.V.S.K.P. and Q.J. performed EMSA experiments; K.V.S.K.P. generated the CG-1 mutants and reporter and effector constructs; A.A.H. cloned the CG-1 mutants into a bacterial expression vector, cloned all mutants into a binary vector, generated transgenic lines and performed transient assays and gene expression analyses. K.V.S.K.P., A.A.H., and A.S.N.R. wrote the manuscript. Q. J. read and provided comments on the manuscript.

## Competing interests

None declared

**Supplementary figure 1. Identification of evolutionarily conserved amino acid residues in the CG-1 domain of six CAMTAs in *Arabidopsis thaliana*.** Clustal W alignment of first 200 amino acids (encompassing NLS and CG-1 domain) sequence in the CG-1 domain of AtCAMTAs 1-6. The stretches of highly conserved amino acid residues in all six CAMTAs proteins are marked by black boxes. Based upon the sequence logo in Fig. 1, the corresponding highly conserved amino acids with >4-bit score are indicated by letters M1-M6 on top of the amino acid.

**Supplementary figure 2. Expression and purification of WT and CG-1 mutants in *E. coli*.** Immunoblot analysis of His-column purified protein of WT and CG-1 mutants (M1 to M6) using anti-His antibody.

**Supplementary figure 3. Significant differences in the binding activity of WT and M5 CG-1 mutant to the DNA probe.**

Comparison of the binding capacity of WT and M5 mutant to *PDF1.4* probe. Based on Fig. 2b and 3b, the signal volume intensity of bound DNA-Protein complex and unbound (DNA probe only) is calculated for each lane at a different protein concentration using image analysis software (Image Lab 6.0.1 version, Bio-Rad) provided by the manufacturer. Lane numbers are indicated on top of the bars.

**Supplementary figure 4. Generation of *RSRE::NOS and mRSRE::NOS* promoter for the reporter construct.**

The sequence of *RSRE* [*CGCGTT*] elements (highlighted in green) included in the upstream region of the *NOS* minimal promoter (highlighted in grey) interspersed by other bases was used for the synthesis of *RSRE::NOS* promoter that was cloned upstream to the luciferase gene. For the synthesis of the mutant version *mRSRE::NOS* promoter, the *RSRE* elements (highlighted as magenta) and surrounding sequences were altered keeping the *NOS* minimal promoter (highlighted as grey) intact.

